# Essential Metabolic Routes as a Way to ESKAPE from Antibiotic Resistance

**DOI:** 10.1101/817015

**Authors:** Angélica Luana C. Barra, Lívia de Oliveira C. Dantas, Luana Galvão Morão, Raíssa F. Gutierrez, Igor Polikarpov, Carsten Wrenger, Alessandro S. Nascimento

**Affiliations:** São Carlos Institute of Physics, University of Sao Paulo. Av. Trabalhador São-carlense, nº 400, São Carlos, SP. Brazil; Department of Parasitology, Institute of Biomedical Sciences, University of Sao Paulo. Av. Prof. Lineu Prestes, 1374 – Cidade Universitária, São Paulo, SP, Brazil

**Keywords:** ESKAPE pathogens, thiamine, pyridoxal 5’-phosphate, antibiotic resistance

## Abstract

The antibiotic resistance is a worldwide concern that requires a concerted action from physicians, patients, governmental agencies and academia to prevent infections and the spread of resistance, to track resistant bacteria, to improve the use of current antibiotics and to develop new antibiotics. Despite the efforts spent so far, the current antibiotics in the market are restricted to only five general targets/pathways highlighting the need for basic research focusing on the discovery and evaluation of new potential targets. Here we interrogate two biosynthetic pathways as potentially druggable pathways in bacteria. The biosynthesis pathway for thiamine (vitamin B1), absent in humans, but found in many bacteria, including organisms in the group of the ESKAPE pathogens (*Enterococcus faecium*, *Staphylococcus aureus*, *Klebsiella pneumoniae*, *Acinetobacter baumanii*, *Pseudomonas aeruginosa* and *Enterobacter* species) and the biosynthesis pathway for pyridoxal 5’-phosphate and its vitamers (vitamin B6), found in *S. aureus*. Using current genomic data, we discuss the possibilities of inhibition of enzymes in the pathway and review the current state of the art in the scientific literature.

## 1 Introduction

Antibiotic resistance is an urgent threat to human health and requires urgent actions from physicians, patients, industries, governmental agencies and the academic community worldwide. According to the last document from the Centers for Disease Control and Prevention (CDC) regarding antibiotic resistance in the United States, from 2013, the number of people with serious infections caused by resistant bacteria reaches two million every year, with at least 23,000 deaths per year directly caused by these infections (1). The situation is similarly warning in Europe, where 25,100 deaths were reported from the European Centre for Disease Prevention and Control in 2007 (2). Globally, 700,000 deaths are estimated every year as a consequence of antibiotic resistance (3). The same CDC document lists four general action lines to address antibiotic resistance: (*i*) preventing infections and the spread of resistance; (*ii*) tracking resistant bacteria; (*iii*) improving the use of current antibiotics; (*iv*) developing of new antibiotics (1).

Since bacteria have a short doubling time and efficient mechanisms for plasmid sharing, the development of antibiotic resistance is a very efficient defense mechanism. So, as previously said by Walsh and Wencewicz, the development of resistance is not a matter of *if*, but rather a matter of *when* (4) and, despite the title of this paper (which is rather provocative), there is no way we can escape from it (5). On the contrary, the development/discovery of new antibiotics will tend to be a continuous goal to be achieved in drug discovery pipelines.

Interestingly, even after what has been named as the ‘golden age’ of antibiotics development, the drugs currently in the market are restricted to only five molecular targets and/or pathways (4): (*i*) the peptidoglycan/cell wall biosynthesis, site of action of beta-lactam antibiotics, for example; (*ii*) the protein biosynthesis, where the ribosome is an important target; (*iii*) DNA replication and RNA transcription; (*iv*) the folate biosynthesis pathway, and (*v*) the disruption of the bacterial membrane. Given the relevance of the antibiotic therapy and the emergence of antibiotic resistance, the available choices of current drugs are very narrow in the context of the mechanism of action restricted to only five molecular/pathway targets.

Antibiotic resistance, as a threat to human health, should be addressed in multiple, and simultaneous ways. For academia, an interesting way to address antibiotic resistance is the discovery and validation of new targets/pathways that could be specifically targeted by new antibiotic candidates. It is worthy of note that, according to Kelly and Davies (3), no new class of antibiotics was discovered and released for routine treatment since the 1980s, highlighting the outstanding role that academia can have in the preliminary research for discovery and validation of druggable targets/pathways.

About 7% of the *E. coli* genome, 303 genes, were shown to be composed of essential genes (5,6). Under stress conditions caused by a limited medium, additional 119 genes show some condition-dependent essentiality, including several genes related to the metabolism and synthesis of essential molecules such as amino acids, and nucleotides (5) and, obviously, some of these targets/pathways can be an interesting choices for the development of new antibiotic candidates. A promising approach to screen compounds for their activity in metabolic pathways was shown by Zlitni and coworkers (7). The authors searched for antibacterial compounds under poor nutrient media and found three potential compounds in this screening strategy (7). Worth of note, the metabolic profile of existing antibiotics showed that the supplementation of vitamins B5, B6, B1, and B2 did not significantly reverse the antimicrobial effect of any of the 24 inhibitors assayed (7), suggesting that the current antibiotics do not explore these essential pathways.

In this context, the enzymatic routes for the biosynthesis of vitamins are very interesting pathways to be explored as potential targets for the discovery of new antibiotic candidates. Vitamin B1, for example, cannot be synthesized by humans, although several microorganisms can synthesize this vitamin, including pathogens. In the absence of thiamin (vitamin B1), the activity of several carbohydrate metabolism enzymes is impaired, including pyruvate dehydrogenase, which connects glycolysis and the citric acid cycle (8). Other thiamin dependent enzymes are transketolase, α-ketoacid decarboxylase, α-ketoacid dehydrogenase and acetolactate synthase (9). A very similar scenario was observed for pyridoxal 5’-phosphate (vitamin B6) in the model Gram-positive organism *Bacillus subtilis* (10).

The pathways involved in the synthesis of vitamins, in particular, vitamin B1 (thiamin) and B6 (pyridoxal), seem to be of great relevance, since they are involved in central processes in the metabolism of carbohydrates and amino acids and the corresponding enzymes are found in most bacteria, fungi and plants but not in humans (8), favoring the development of specific drugs with minimal side effects due to the interaction with the host. However, a few questions still stand: (i) how feasible are the targets involved in the pathways for the biosynthesis of thiamin and pyridoxal for the microorganisms with a higher emergency in terms of resistance, or ESKAPE: *Enterococcus faecium, Staphylococcus aureus, Klebsiella pneumoniae, Acinetobacter baumanii, Pseudomonas aeruginosa* and *Enterobacter* species? (ii) What enzymes are present and what do we know about them?

Here we used the available genomic and proteomic data to address these questions focusing on the ESKAPE pathogens. Vitamins B1 and B6 were chosen for this analysis, since the biosynthetic pathways for these enzymes have already been validated as molecular targets for some human pathogens, such as thiamin for *Plasmodium falciparum* (11), or pyridoxal for *P. falciparum* (12) or *Trypanosoma brucei* (13), for example.

## 2 Methods

The analyses provided here are the result of the interrogation of the enzymes in the biosynthesis pathways for thiamin and pyridoxine phosphate or pyridoxal phosphate using the KEGG database (14,15). For this purpose, the KEGG pathway for thiamin metabolism (map 00730) was listed for the ESKAPE pathogens using, whenever possible, commercial strains rather than specific or antibiotic-resistant strains. Briefly, for each pathogen (*Enterococcus faecium*, *Staphylococcus aureus*, *Klebsiella pneumoniae*, *Acinetobacter baumanii*, *Pseudomonas aeruginosa* and *Enterobacter)* a KEGG search for the maps of thiamin and pyridoxal metabolism was done, looking for the existing genes in commercial strains for each of the ESKAPE organisms. For the sake of comparison, some additional resistant strains were listed, with no differences in the observed genes for the pathways under study. A similar search was carried out for the vitamin B6 metabolism (map 00750) in KEGG, interrogating the existing enzymes for the ESKAPE pathogens in comparison with humans. The analysis was focused in the enzymes in the biosynthesis of B1 and B6 vitamin in an attempt to identify the most promising targets for future medicinal chemistry studies. Finally, a table listing the existing genes for each pathogen was compiled and is presented in the following sections.

Using the existing literature, we sought to identify whenever possible the cases of enzymes with functional redundancy with another enzyme for the same organism. The identified cases were discussed as less promising targets for medicinal chemistry campaigns. Additionally, the existing data for enzyme inhibition using natural or synthetic compounds was compiled to provide initial proof-of-concept clues about the *druggability* of the identified promising targets.

When necessary, the sequence of an enzyme of the pathway was used to search for homologues using BLAST (16) searches against the PDB (17) or the non-redundant database of proteins using BLAST default parameters, i.e., minimum expected threshold of 10, a word size of 6 and the BLOSUM62 substitution matrix.

## 3 Results and Discussion

### 3.1 Thiamin Biosynthesis

The biosynthesis pathway for thiamin involves two branches, as shown in Figure 1. In summary, thiamin is synthesized from 4-amino-5-hydroxymethyl-2-methylpyrimidine (HMP, superior branch in Figure 1) and 5-(2-hydroxyethyl)-4-methylthiazole (THZ, inferior branch). Both compounds are phosphorylated by ThiD and ThiM, respectively. Finally, the enzyme ThiE, central to the pathway, is responsible to synthesize thiamine phosphate (TMP) by merging the two branches (18). Thiamin phosphate can be dephosphorylated to thiamine and then pyrophosphorylated to thiamin diphosphate (TPP) by TPK. Alternatively, thiamin can be degraded to HMP and THZ by TenA, a thiaminase II enzyme.

According to the KEGG pathway database, humans lack many of the enzymes of this pathway, but not all of them. In humans, a TPK (TPK1) enzyme is found, with UNIPROT ID Q9H3S4. Other than that, the remaining enzymes in the biosynthetic pathway for thiamin are missing in humans, making them very attractive for the design of chemical probes that could be used as proof-of-concept compounds.

The analysis of the thiamin pathway for *Enterococcus faecium* (ATCC 8459) in the KEGG database did not identify genes for ThiE, ThiM and TenA. For this species, only ThiD and TPK were positively identified. However, a BLAST search for TenA homologous within the non-redundant database restricted to *Enterococcus faecium* (TAXID 1352) identified a single result for thiaminase II (GenBank: SAM74984.1), four results for ThiE (SAZ10134.1, WP_086323306.1, WP_010732763.1 and WP_072538983.1) and two results for ThiM (SAZ10238.1, EJY48288.1). The small number of hits in the BLAST search may suggest some issues in the annotation.

Interestingly, when the *Enterococcus faecalis* thiamin pathway is compared with the pathway observed for *E. faecium*, many differences are found. The KEGG pathway for *E. faecalis* (ATCC 29212) shows that the entire pathway, as described in Figure 1 is found: ThiD, ThiM, ThiE, TPK and TenA, in contrast with *E. faecium*. Additionally, there is no change in the repertoire of enzymes in the pathway for the Vancomycin-resistant strain V583 as compared to the ATCC 29121 strain.

For *Staphylococcus aureus* (NCTC 8325) and *Klebsiella pneumoniae* (subsp. pneumoniae ATCC 43816 KPPR1), the KEGG pathway indicates that all the enzymes in the pathway are observed, with no changes to an *S. aureus* resistant strain such as COL (MRSA) as compared to the NCTC 8325. For *Acinetobacter baumannii* (ATCC 14978), all enzymes are observed but TPK. Instead, thiamin phosphate may be converted directly to thiamin diphosphate by a thiamin-monophosphate kinase, and then converted to thiamin triphosphate by an adenylate kinase. In the case of *Pseudomonas aeruginosa* (NCGM 1900), some enzymes are missing: ThiD, TenA, TPK. Finally, *Enterobacter sp.* (638) has ThiD, ThiE, ThiM and misses TenA and TPK.

The overall panel of enzymes for the thiamin pathway for the ESKAPE pathogens is summarized in Table 1.

**Table 1.**
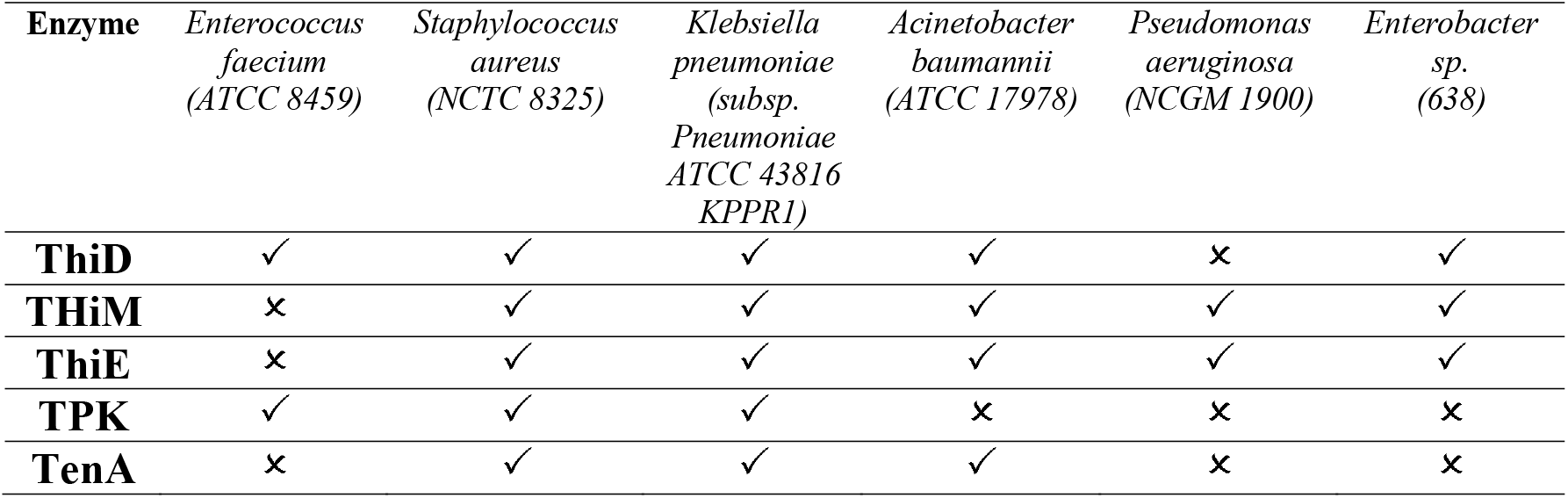
Panel of observed enzymes for the thiamine pathway in ESKAPE pathogens.

TenA was shown to play a dual role in thiamine synthesis and salvage (19): beyond its function to hydrolyze thiamin, TenA also deaminates aminopyrimidine to form 4-amino-5-hydroxymethyl-2-methylpyrimidine (HMP), with the latter activity being about 100 times faster than the former (19,20). In some organisms, such as *Bacillus subtilis*, the deaminase activity is somewhat redundant with the ThiC (ThiA) activity, since both activities provide HMP or HMP phosphate, that can be further phosphorylated by ThiD (9).

In the other branch of the thiamin biosynthesis pathway, thiazole phosphate can be generated from thiazole alcohol incorporated from the medium and phosphorylated by ThiM or it can be synthesized in a series of enzymatically catalyzed reactions using amino acids such as glycine or tyrosine as substrates (9). In this context, according to Begley and coworkers (9), lesions in the ThiM gene were observed to be prototrophic. In contrast, selective mutations of ThiD and ThiE led to the requirement of externally provided thiamin (9).

ThiE is central to the pathway since the enzyme merges the two branches to finally synthesize thiamin phosphate (Figure 1). It is important to highlight that many of the mutational analysis was carried out in *E. coli* and, as Table 1 shows, there is significant variance among bacterial species.

Taking together, the analysis of the literature suggests that ThiD and ThiE may represent interesting targets for the development of chemical probes to further evaluate their function. TPK is expressed in humans and the inhibition of this enzyme could lead to harmful effects on the human host. TenA inhibition could be overcome by the somewhat functional redundancy with thiC (thiA) and ThiM function can be dispensable since thiazole phosphate can be synthesized *de novo*.

ThiD and ThiE are found in most ESKAPE organisms. Exceptions are *P. aeruginosa*, which lacks ThiD and *E. faecium* that lacks ThiE, according to KEGG data. For ThiE, in particular, it was reported that 4-amino-2-trifluoromethyl-5-hydromethylpyrimidine (CF_3_-HMP), an HMP analogue, can be converted by ThiD to CF_3_-HMP pyrophosphate, which in turn inhibits ThiE (8,21). This enzymatic inhibition culminated with the inhibition of *E. coli* growth (8), suggesting that this strategy might be promising for the development of new inhibitors.

In terms of the structural biology, some crystal structures of ThiD are available by the time of the writing of this paper, including the enzymes from *Salmonella enterica* (PDB ID 1JXH (22)), *Clostridioides difficile* (PDB ID 4JJP), *B. subtilis* (PDB ID 2I5B (23)), *Thermus thermophilus* (PDB ID 1UB0), *A. baumannii* (PDB ID 4YL5), *Bacteroides thetaiotaomicron* (PDB ID 3MBH) and the bifunctional enzyme from *Saccharomyces cerevisiae* (PDB ID 3RM5). Typically, ThiDs are folded as typical ribokinases, with a central 8-stranded β-sheet surrounded by 8 α-helices and structural studies suggest that some surface loops may have a structural change based on the presence of the nucleotide (23).

Interestingly, the *S. aureus* pyridoxal kinase enzyme (*Sa*PdxK) has a dual role, phosphorylating pyridoxal and pyridoxine in the pyridoxal *de novo* biosynthesis pathway as well as HMP in the thiamin biosynthesis pathway, with a K_M_ almost 20 times greater for HMP than for pyridoxal and k_cat_ values 3 times faster for pyridoxal (24). So, SaPdxK has some redundancy with *S. aureus* ThiD (SaThiD), with less efficiency, though.

*Sa*ThiD was shown to be potentially inhibited by Rugulactone (Ru0), a natural product as well as by its derivatives Ru1 and Ru2, with IC_50_ values ranging from 14 to 32 μM (25). In the absence of thiamine in the medium, the MIC observed for Ru1 was four times lower than in the presence of thiamine for *L. monocytogenes* (25), suggesting that the inhibitory effect of Rugulactone is due to ThiD inhibition, although Rugulactone also inhibits other kinases.

For ThiE, the crystal structures of a few enzymes are available, including the enzyme from *Pyrococcus furiosus* (PDB ID 1XI3), *B. subtilis* (PDB IDs 1G4T, 3O15, 3O16, 1G69, 1G4E, 1G67, 1G4P, 1YAD (26)), *Mycobacterium tuberculosis* (3O63) and for the bifunctional enzyme from *Candida glabrata* (PDB IDs 3NL2 and 3NM1 (27)). No crystal structure of an ESKAPE pathogen ThiE is available to date. From the structural point of view, ThiE is typically folded as an α/β TIM barrel, with thiamine binding at the top of the barrel, as elegantly shown in the set of crystal structures determined for the *B. subtilis* enzyme (28).

At this point, an important question has to be addressed: can the blockade of thiamine *de novo* biosynthesis pathway be overcome by internalization of the vitamin? In principle, several microorganisms can import thiamine from the environment (29). So, how effective can be the inhibition of ThiD and ThiE particularly for pathogenic bacteria? This question can be addressed in several levels. On the first level, the essentiality of ThiD, for example, has been shown some pathogenic microorganisms (23), such as *Streptococcus pneumoniae* (30), *Haemophilus influenzae* (31) and *Mycobacterium tuberculosis* (32). This is not the case for other bacteria. For *B. subtilis*, for example, ThiD gene was shown to be dispensable (23). Whether ThiD and ThiE are essential for the ESKAPE pathogens is still an open question to be addressed by further investigation. On a second level, Nodwell and coworkers showed that *Listeria monocytogenes*, which is known to uptake thiamine from the environment, is still sensible to Rugulactone and its derivatives Ru1 and Ru2 (25). When grown in a chemical defined media without thiamine, the inhibitory effects of Rugulactone are potentiated with a 4-fold reduction in the MIC (25). However, the inhibitory effect observed even in the presence of thiamine highlight that the blockade of the *de novo* biosynthesis pathway is still a promising strategy. As a third level, the natural compound bacimethrin, isolated from *Bacillus megaterium* and from *Streptomyces albus* is known to be toxic for bacteria with MIC at low micromolar range (33,34). The mechanism of action of bacimethrin is based on the formation of 2’-methoxy-thiamine where bacimethrin is used by ThiD instead of HMP. 2’-methoxy-thiamine pyrophosphate inhibits *E. coli* growth at concentrations 15 times lower than bacimethrin (35). Interestingly, some *Salmonella enterica* thiD mutants were shown to be bacimethrin resistant (36). Together, these data suggest that a very promising approach can be devised by the design of suicide drugs, i.e., compounds that are recognized by the enzymes in the pathway but lead to final compounds that can’t be used as thiamine substituents. Bacimethrin is a good example of a suicide inhibitor and, although its effects can be reverted by increased thiamine concentrations, the vitamin concentration that pathogens are exposed are usually defined within a narrow range.

### 3.2 Pyridoxal Biosynthesis

Pyridoxal, pyridoxine and pyridoxamine, together with their respective phosphate esters compose the six vitamers for vitamin B6 and can be interconverted (37). There are two biosynthetic routes for pyridoxal 5’-phosphate: a deoxyxylulose 5-phosphate-dependent pathway found in some bacteria and a deoxyxylulose 5-phosphate-independent pathway found in all kingdoms (37,38). This second and widespread pathway, curiously, depends on only two enzymes, as shown in Figure 2.

In this pathway, the enzyme Pdx2 converts L-glutamine into L-glutamate and an ammonium molecule. The latter is thought to diffuse to Pdx1 active site, which also uses glyceraldehyde 3-phosphate and D-ribose 5-phosphate to synthesize pyridoxal 5’-phosphate (37,39). Interestingly, to properly synthesize pyridoxal 5’-phosphate, a complex involving twelve Pdx1 molecules and twelve Pdx2 molecules has to be assembled (40–42) and this assembly is further stabilized by the interaction with L-glutamine (43).

Similar to what is observed to the thiamine biosynthesis pathway, humans lack the genes for Pdx1 and Pdx2, making the development of specific antibiotic candidates an attracting strategy. In the same direction, for some pathogenic organisms such as *Helicobacter pylori* (44), *Mycobacterium tuberculosis* (45), and *Streptococcus pneumoniae* (46), the depletion in vitamin B6 resulted in reduced virulence, indicating that the biosynthesis of this vitamin may be a good strategy for the design of new antibiotic candidates.

Very interestingly and in contrast to what was observed for the thiamine pathways, from the ESKAPE pathogens, *S. aureus* is the only microorganism that has the ribose 5-phosphate dependent pathway for the biosynthesis of pyridoxal 5’-phosphate, as shown in Table 2. A single enzyme (Pdx1 or Pdx2 alone) is never observed for these organisms (Table 2).

**Table 2.** Panel of observed enzymes for the pyridoxal 5’-phosphate pathway in ESKAPE pathogens.

| Enzyme | *Enterococcus*<br>*faecium*<br>(ATCC 8459) | *Staphylococcus*<br>*aureus*<br>(NCTC 8325) | *Klebsiella*<br>*pneumoniae*<br>(subsp.<br>*Pneumoniae*<br>ATCC 43816<br>KPPR1) | *Acinetobacter*<br>*baumannii*<br>(ATCC 17978) | *Pseudomonas*<br>*aeruginosa*<br>(NCGM 1900) | *Enterobacter*<br>*sp.*<br>(638) |
| --- | --- | --- | --- | --- | --- | --- |
| **Pdx1** | × | ✓ | × | × | × | × |
| **Pdx2** | × | ✓ | × | × | × | × |

A comparison between the pyridoxal/pyridoxine phosphate pathways for *E. faecium* and *E. faecalis* shows that both pathogens miss the Pdx enzymes. On the other hand, some other pathogenic organisms, including organisms with high antibiotic resistance rates, have already been identified as susceptible to modulations of the pyridoxal phosphate biosynthesis pathway. Some organisms which have been the focus of basic research include *Plasmodium falciparum* and *Plasmodium vivax*, the malaria pathogens (47–50).

In terms of the structural biology of the enzymes in the vitamin B6 biosynthesis pathway, the *S. aureus* Pdx1 enzyme (UNIPROT ID Q2G0Q1) is a close homologue of *Bacillus subtilis* Pdx1 (81% identity, PDB ID 2NV1 (42)), *Geobacillus stearothermophilus* Pdx1 (78% identity, PDB ID 1ZNN (51)) and *Thermus thermophilus* enzyme (67% identity, PDB ID 2ZBT). *S. aureus* Pdx2, has *Geobacillus kaustophilus* (PDB ID 4WXY (40)), *Geobacillus stearothermophilus* (PDB ID 1Q7R) and *B. subtillis* (PDB IDs 2NV0 and 2NV2 (42)) enzymes as its close homologues, with 60%, 60% and 58% identity in sequence similarity respectively.

In terms of its structure, the Pdx1 enzyme has a typical (β/α)_8_ barrel fold with a central 8-stranded β-barrel surrounded by 8 α-helices (51). The most impressing feature in the Pdx1 enzyme structure is its quaternary arrangement, where six Pdx1 enzymes interact with each other to form a ‘*donut-like*’ arrangement with about 100 Å in diameter. For Pdx2, a Rossman fold is observed with an α/β/α sandwich topology (42). Curiously, in the active PLP synthase complex, 12 Pdx1 molecules (a two-layer *donut*) interact with 12 Pdx2 enzymes. However, Pdx2 molecules do not interact with each other in this complex (42).

Using a structural homology model, Reeksting and coworkers identified some ribose 5’-phosphate analogues with interesting *in vitro* inhibitory effects on the enzyme Pdx1 from *P. falciparum* (12). The compounds were identified in a structure-based computational screening campaign and were shown to have inhibitory effects *in vivo*, as well as the *in vitro* effect. The authors also showed that the *in vivo* effects could be at least partially suppressed in a mutant strain overexpressing Pdx1 and Pdx2, in a clear indication that the effect of the analogues was due their inhibition of the pyridoxal 5’-phosphate biosynthesis pathway (12).

The inhibition of the Pdx2 activity by the glutamine analogue acivicin was also demonstrated to be a feasible strategy for the inhibition of the vitamin B6 biosynthesis pathway (52). Raschle and coworkers showed that when inhibited by acivicin, Pdx2 is incapable of interacting with Pdx1, thus disrupting the pyridoxal 5’-phosphate synthase activity. Interestingly, acivicin is a covalent inhibitor, that binds to a cysteine residue. So, it seems that multiple strategies may be available for the design of new binders, including the inhibition of Pdx2 (covalent and non-covalent), the inhibition of Pdx1 and possibly the inhibition of the assembly of the 24-mer complex with Pdx1/Pdx2. Another possibility to address the inhibition of PLP biosynthesis might be at the level of the Pdx1 transcription, which was shown to be regulated by PdxR (46). In principle, if PdxR action could be regulated by an exogenous chemical probe, the PLP synthase activity would be regulated as well. However, a proof-of-concept experimental validation is still necessary.

Finally, a salvage pathway for vitamin B6 is found in many bacteria, as well as in humans. In *E. coli*, it involves two enzymes: PdxK and PdxY. These kinases act phosphorylating the vitamers pyridoxal, pyridoxine and pyridoxamine into their phosphorylated forms (23,37,53). Interestingly, in some organisms, including pathogenic microorganisms such as *S. aureus*, the PdxK activity is carried out by the HMP kinase ThiD, showing some functional convergence between the vitamin B6 and B1 pathways (24). Not surprisingly, this enzyme has also been shown to be validated druggable target for some pathogens, such as *Trypanosoma brucei* for example (13), although specificity may be an important issue, since humans also have genes for PdxK known to be inhibited by drugs like theophylline, for example, with known neurotoxic effects (54).

Again, the same question asked for the thiamine biosynthesis pathway can be asked here: how can the microbial uptake of vitamin B6 overcome the blockade of the biosynthetic pathway? Some evidences suggest that the blockade might be effective. For example, in *S. pneumoniae*, the deletion of Pdx1 resulted in defective growth of the bacteria. The growth was shown to be restored under increased concentrations of vitamin B6 (46). A similar scenario was also observed for *M. tuberculosis* (45), suggesting that the uptake might not be enough to meet the pathogen’s requirement of the vitamin.

In conclusion, the biosynthesis pathways for vitamin B1 (thiamine) and B6 (pyridoxal 5’-phosphate) may be interesting molecular targets for the development of new chemical probes aiming to inhibit the synthesis of these essential cofactors in pathogenic organisms. Model compounds, mainly based on substrate analogues showed that inhibition of the vitamin B1 biosynthetic enzymes, ThiD and ThiE, as well as Pdx1, Pdx2 and PdxK in pyridoxal 5’-phosphate biosynthesis pathway have shown an initial proof of concept for the model and it is up the scientific community and/or researcher in industry to explore these target further.

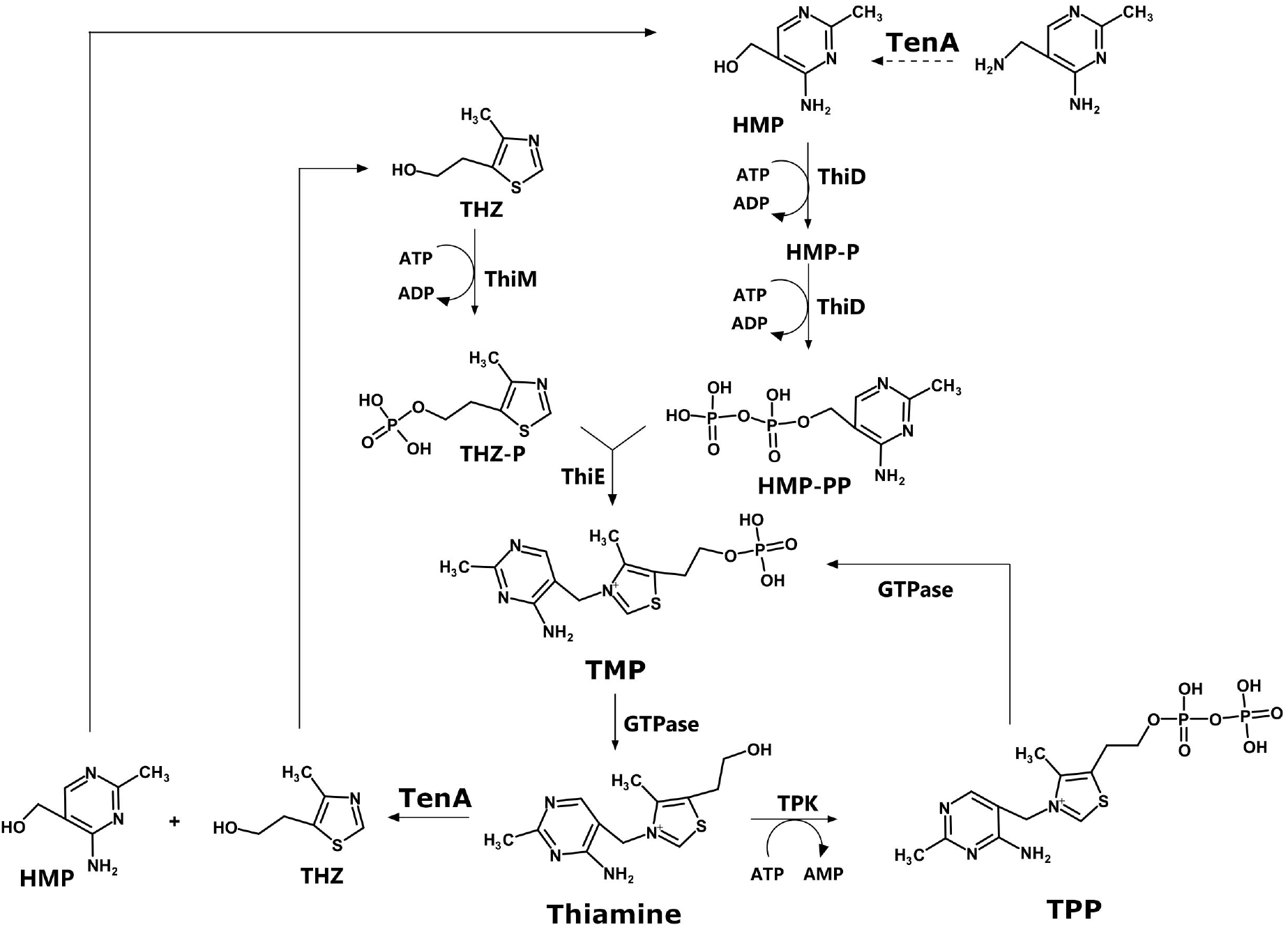

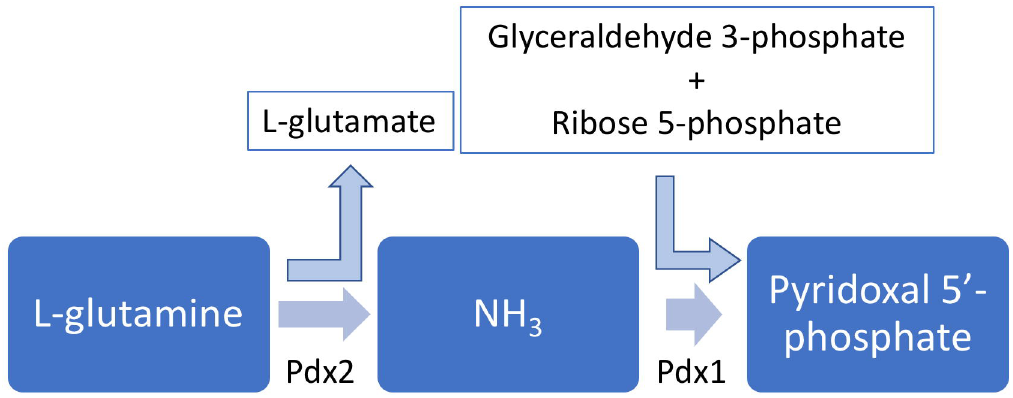

## 4 Conflict of Interest

The authors declare that the research was conducted in the absence of any commercial or financial relationships that could be construed as a potential conflict of interest.

## 5 Author Contributions

CW, IP and ASN conceived the project. LGM and RFG analyzed the thiamine pathway, while ALCB and LOCD analyzed the pyridoxal pathway. All authors wrote and approved the manuscript.

## 6 Funding

Financial support was provided by Fundação de Amparo à Pesquisa do Estado de São Paulo (FAPESP) through grants 2017/18173-0, 2015/26722-8 and 2015/13684-0 and by Conselho Nacional de Desenvolvimento Científico e Tecnológico (CNPq), through grants 303165/2018-9 and 406936/2017-0.

## 7 Acknowledgments

The authors thank the funding agencies FAPESP and CNPq. We are also indebted to Maria Auxiliadora M. Santos, Lívia Regina M. Margarido, Josimar Sartori and João Fernando Possatto for their technical support. This manuscript has been released as a Pre-Print at BioRxiv (55).

## Notes

#### Summary of Updates

The manuscript was extended to cover some additional discussion about the potential inhibition of ThiD and ThiE. Additionally, a correction was introduced to avoid a misunderstanding between S. aureus PdxK and ThiD.

